# In *Mycobacterium abscessus*, the stringent factor Rel regulates metabolism, but is not the only (p)ppGpp synthase

**DOI:** 10.1101/2020.11.07.372714

**Authors:** Augusto César Hunt-Serracín, Misha I. Kazi, Joseph M. Boll, Cara C. Boutte

**Affiliations:** Department of Biology, University of Texas, Arlington

## Abstract

The stringent response is a broadly conserved stress response system that exhibits functional variability across bacterial clades. Here, we characterize the role of the stringent factor Rel in the non-tuberculous mycobacterial pathogen, *Mycobacterium abscessus* (*Mab*). We found that deletion of *rel* does not ablate (p)ppGpp synthesis, and that *rel* does not provide a survival advantage in several stress conditions, or in antibiotic treatment. Transcriptional data show that Rel*_Mab_* is involved in regulating expression of anabolism and growth genes in stationary phase. However, it does not activate transcription of stress response or antibiotic resistance genes, and actually represses transcription of many antibiotic resistance genes. This work shows that there is an unannotated (p)ppGpp synthetase in *Mab*.

**Importance:** In this study, we examined the functional roles of the stringent factor Rel in *Mycobacterium abscessus* (*Mab*). In most species, stringent factors synthesize the alarmone (p)ppGpp, which globally alters transcription to promote growth arrest and survival under stress and in antibiotic treatment. Our work shows that in *Mab*, an emerging pathogen which is resistant to many antibiotics, the stringent factor Rel is not solely responsible for synthesizing (p)ppGpp. We find that Rel*_Mab_* downregulates many metabolic genes under stress, but does not upregulate stress response genes and does not promote antibiotic tolerance. This study implies that there is another critical but unannotated (p)ppGpp synthetase in *Mab*, and suggests that Rel*_Mab_* inhibitors are unlikely to sensitize *Mab* infections to antibiotic treatment.

## Introduction

Bacteria must adjust their physiology to permit survival in fluctuating conditions. The stringent response is a conserved signaling system that promotes survival of many species in stress and antibiotics by altering the transcription of about a quarter of the genome (1–5). In this work, we profile the role of Rel, the sole annotated stringent factor, in the non-tuberculous, rapidly-growing *Mycobacterium abscessus* (*Mab*). *Mab* is an opportunistic pathogen that both lives in the environment, and causes skin and respiratory infections which are increasingly prevalent in Cystic Fibrosis patients (6). *Mab* infections are difficult to treat due to intrinsic resistance to many antibiotics (7), and high tolerance under stress to almost all antibiotics tested (8, 9). One proposed strategy to help treat such antibiotic-recalcitrant infections is to inhibit regulatory systems, like the stringent response, which promotes antibiotic tolerance (10–13).

The conserved aspect of the stringent response is the synthesis, upon stress, of the hyperphosphorylated guanine (p)ppGpp. Once made, (p)ppGpp affects transcription in different ways (14–17) and also directly modulates replication (18, 19), nucleotide metabolism (20–22), ribosome maturation (23, 24) and translation (25–27).

There are a handful of different protein families that synthesize (p)ppGpp across bacterial clades. The most widely conserved are the Rel/Spo Homolog or RSH proteins, which contain, from N-terminus to C-terminus, a (p)ppGpp hydrolase domain, a (p)ppGpp synthase domain, and regulatory TGS and ACT domains (28). Some RSH proteins associate with the ribosome and sense amino acid starvation by detecting when ribosomes have stalled due to an uncharged tRNA being in the A site (29, 30). Some RSH proteins detect other types of stress or nutrient deprivation via other mechanisms (31–34). Many species have only one RSH-type protein, which is competent for both (p)ppGpp hydrolysis and synthesis (28). Many species encode Small Alarmose Synthetase (SAS) and Small Alarmone Hydrolase (SAH) proteins, which contain only (p)ppGpp synthetase and hydrolase domains respectively (30, 35–38). Because SAS proteins do not have regulatory domains, they are typically controlled transcriptionally (30). There are a few other (p)ppGpp synthesizing domains that have been preliminarily studied. ToxSAS proteins are part of toxin-antitoxin systems and synthesize (p)ppGpp as a toxin - these are found in both phage and bacterial genomes (39). *Streptomyces antibioticus* has a polynucleotide phosphorylase (PNPase) which can also synthesize (p)ppGpp (40–43).

The physiological outputs of the stringent response vary across species, but there are conserved themes. First, the stringent response generally downregulates genes required for growth, such as ribosome and cell wall synthesis factors, and it alters transcription of central metabolism to prioritize survival rather than construction of new cells (2, 3, 17, 44). Stringent inhibition of growth (44–50) indirectly protects against some stresses and antibiotics that interfere with growth factors. In many species, the stringent response upregulates stress response genes such as heat shock proteins, hibernation factors, and stress-specific transcription factors (3, 51, 52) and promotes survival in stress (13). The stringent response also helps many bacteria survive through antibiotic treatment by promoting antibiotic tolerance (12).

The stringent response has been studied in *Mycobacterium tuberculosis* (*Mtb*) and *Mycobacterium smegmatis* (*Msmeg*). *Mtb* has one major (p)ppGpp synthetase, Rel*_Mtb_* which makes (p)ppGpp when respiration is inhibited, in stationary phase, and in total carbon and nitrogen starvation (46, 53). Rel*_Mtb_* also promotes survival of *Mtb* during nutrient and oxygen starvation, stationary phase (46) and chronic infection of mice (1) and guinea pigs (54, 55). Importantly, Rel*_Mtb_* also makes *Mtb* more tolerant to the first-line clinical antibiotic isoniazid during starvation and infection in mice (11). A PNPase enzyme in *Mtb*, Rv2783, has been shown to have weak (p)ppGpp synthetase activity (56), and the *Mtb* genome contains another gene with a predicted GTP pyrophosphokinase domain, Rv1366. However, the Δ*rel_Mtb_* strain does not synthesize measurable (p)ppGpp in starvation conditions (46), so these other potential synthetases are either inactive or not activated in the conditions tested.

*Msmeg* has an RSH protein, Rel*_Msmeg_*, which can both synthesize and hydrolyze (p)ppGpp (57–60). Although the Δ*rel_Msmeg_* strain showed defects in biofilm formation and stationary phase viability, it can still synthesize (p)ppGpp (57, 61). The secondary (p)ppGpp synthetase, RelZ*_Msmeg_*, has an RNaseHII domain in addition to the conserved SAS ppGpp synthetase domain (59), and also synthesizes pGpp (62). Strains missing both synthetases have further biofilm and aggregation defects, but can still synthesize some (p)ppGpp (62), indicating that there is a third, uncharacterized synthetase in *Msmeg*.

In this study we examined the Δ*rel* strain of *Mab*. Rel*_Mab_* is an RSH protein. There are no other RSH genes, and no SAS genes in the *Mab* genome. We found that the Δ*rel_Mab_* strain still makes (p)ppGpp. The Δ*rel_Mab_* strain does not exhibit survival defects in several stress conditions including antibiotic treatment, but has a growth defect relative to wild type. We measured transcriptional changes in Δ*rel_Mab_* relative to wild type and found it helps downregulate many metabolic pathways in stationary phase.

## Results

In order to explore the role of *rel_Mab_*, we built a strain of *Mab* ATCC19977 with a deletion of the *rel* gene (MAB_2876), which has the canonical RSH gene structure, including a (p)ppGpp hydrolase domain, a (p)ppGpp synthetase domain, a TGS domain and an ACT domain. We measured (p)ppGpp in both the wild-type *Mab* and Δ*rel_Mab_* strains, in logarithmic phase and carbon starvation (Fig. 1). We found that both the wild-type and Δ*rel_Mab_* strain produce (p)ppGpp in both log. phase and starvation, and the amount of ppGpp is increased in both strains in starvation.

**Figure 1.**
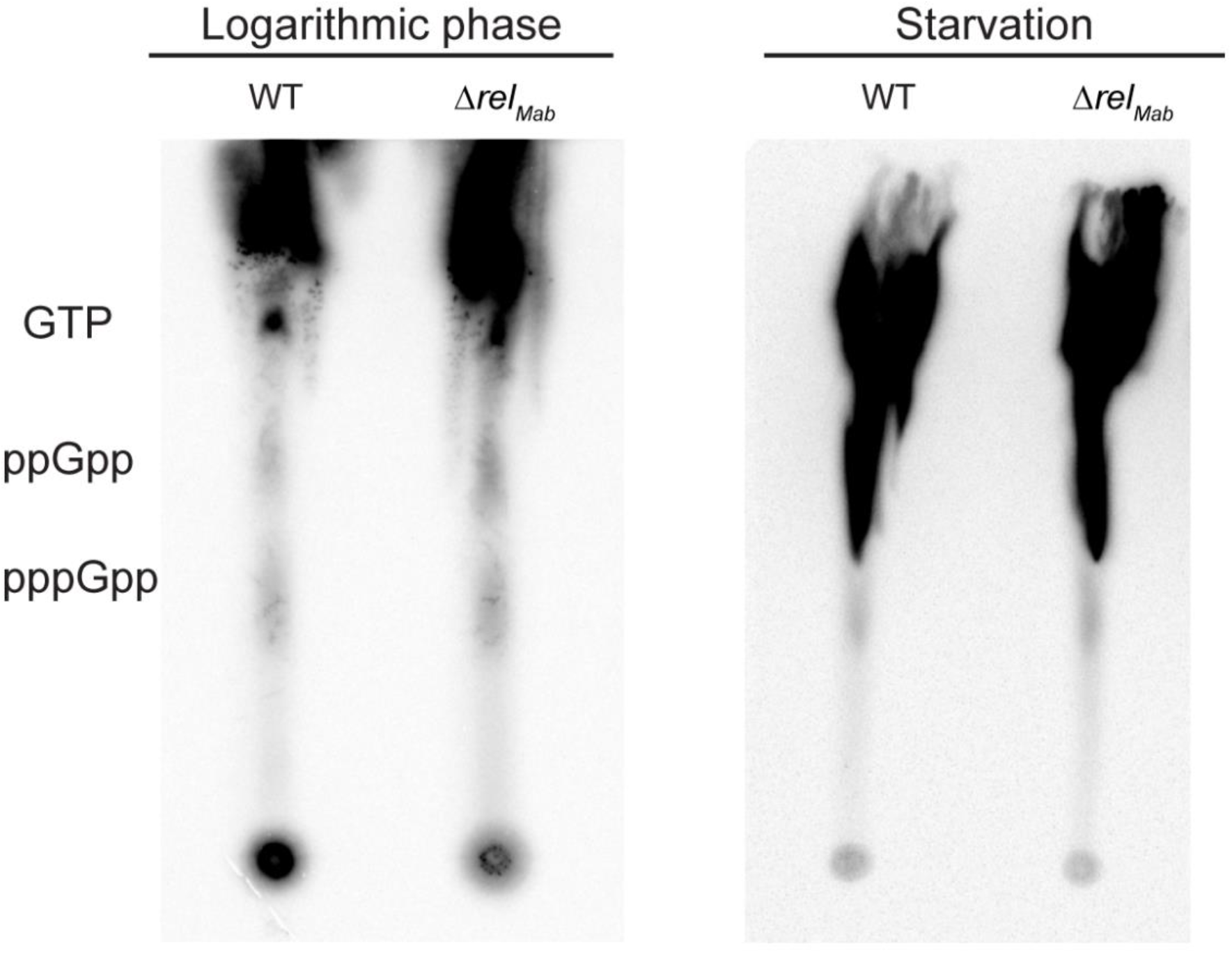
Thin-layer chromatography (TLC) of nucleotides extracted from *M*. *abscessus* strains. Guanine nucleotides extracted from ^32^P-labeled *M. abscessus* strains from logarithmic phase and starvation.

In many species, *rel* orthologues promote survival during stress conditions, such as stationary phase, acid stress, starvation, or oxidative stress (5, 35, 46, 49, 63, 64). To evaluate the physiological role of *rel* in *Mab*, we assayed survival of wild-type, Δ*rel_Mab_* and complemented strains upon and after transfer to either carbon starvation (Fig. 2A), salt stress (Fig. 2B), oxidative stress, (Fig. 2C) or acidic media (Fig. 2D). Treatment with these stressors did not induce measurable differences in growth or survival of Δ*rel_Mab_* relative to wild-type and the complemented strain. Thus, Rel*_Mab_* does not regulate responses to these stresses under the conditions tested, or at least not enough to affect growth or survival. We also found that Rel*_Mab_* does not promote survival in stationary phase (Fig. 3A). Robust survival in the Δ*rel_Mab_* in all stress conditions is likely aided by the continued synthesis of (p)ppGpp under stress (Fig. 1). Growth curves showed that Δ*rel_Mab_* had a significant growth defect relative to wild type and complemented strains (Fig. 3B).

**Figure 2.**
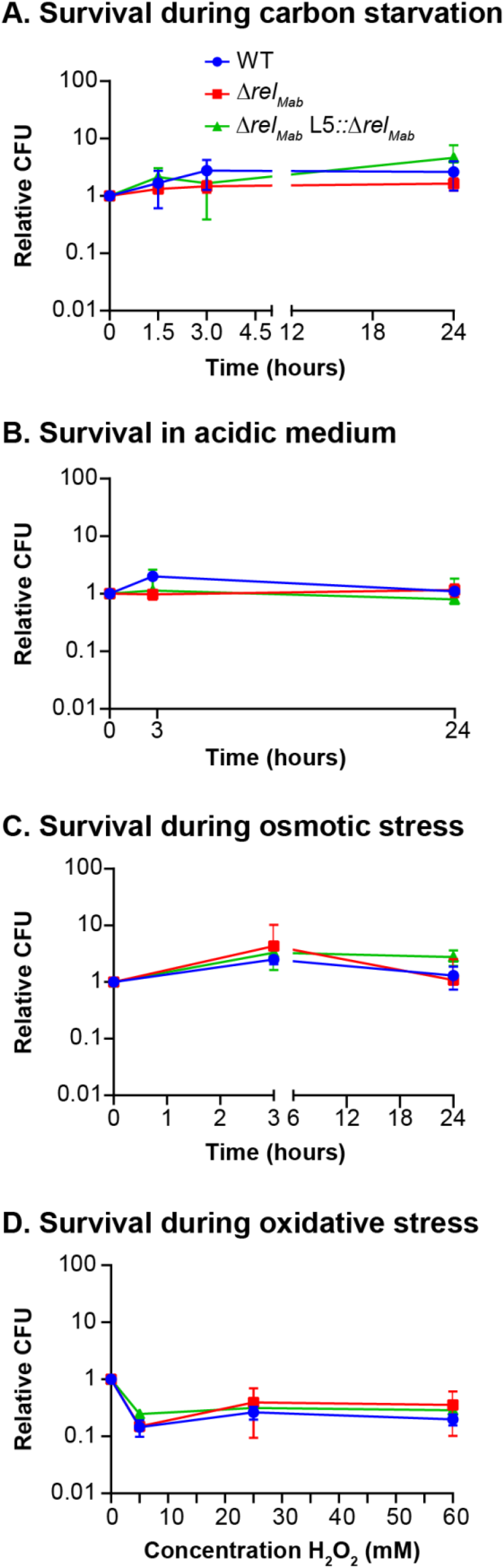
Contribution of *rel_Mab_* to survival in various stresses. (A) Colony-forming units (CFU) of wild type *Mycobacterium abscessus* strain ATCC19977 (blue), Δ*rel_Mab_* (red), and the complemented strain **Δ***rel_Mab_* L5::*rel_Mab_* (green) in Hartman’s du Bont medium with no glycerol and Tyloxapol as a detergent. (B) CFU in 7H9 Middlebrook medium with a pH of 4. (C) CFU in Lennox LB with 1M of NaCl. (D) CFU in Hartmans du Bont medium with 5mM, 25mM or 60mM of tert-butyl peroxide after 24 hours. Relative CFU is calculated by taking the ratio between each CFU value and the initial CFU value at time zero. All data points are an average of three biological replicates. Error bars represent standard deviation. There are no significant differences in any of these data by a two-tailed student’s t-test.

**Figure 3.**
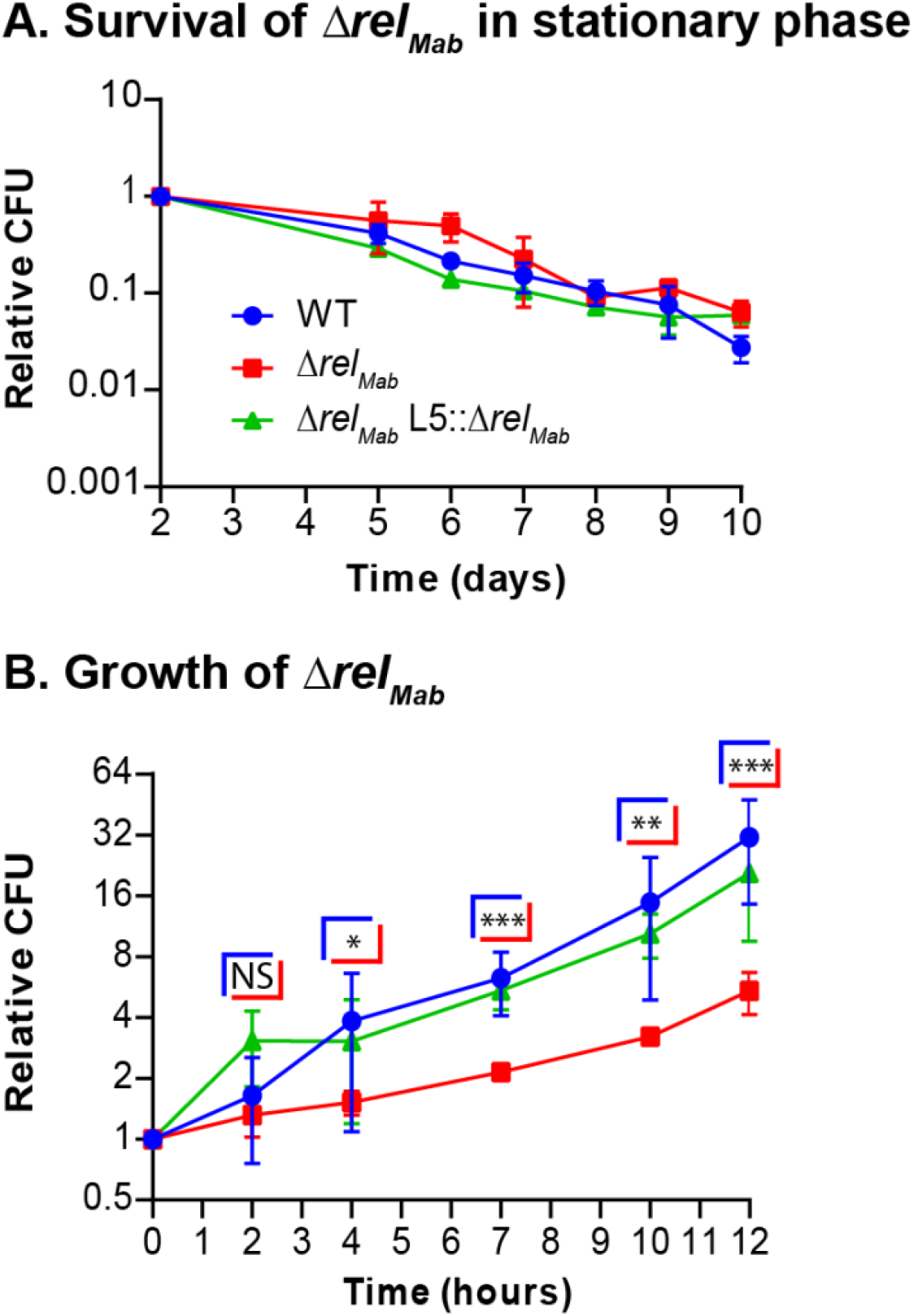
Growth and stationary phase survival of Δ*rel_Mab_*. (A) Colony-forming units (CFU) in stationary phase in 7H9 media. The 2 day time point is 48 hours after diluting logarithmic phase cultures to OD_600_=0.05. (B) CFU during logarithmic phase growth in 7H9 medium. Graph is set at a log2 scale. Doubling-times ± 95% confidence interval = wild type 2.15 h ± 0.08; Δ*rel_Mab_* 4.67 h ± 4.26; Δ*rel_Mab_* L5::*rel_Mab_*; Δ*rel_Mab_* L5::*rel_Mab_* 2.84 ± 1.85. *P*-values are for the wild-type compared to Δ*rel_Mab_. P*-values for *t*=4, *P* = 0.0415; *t*=7, *P* = 0.00015; *t*=10, *P* =0.006; *t*=12, *P* =0.00029. *P*-values between wild-type and the complemented strain were not significant. *P-* values between strains in stationary phase were not significant (data not shown). Asterisks represent significance as measured by the two-tailed student’s *t*-test; * = *P* ≤ 0.05; ** = *P* ≤ 0.01; *** = *P* ≤ 0.001; n.s. = *P* > 0.05.

Because the stringent response is a major activator of antibiotic tolerance and persistence in many species (5, 65, 66), we sought to assess how Rel contributes to antibiotic tolerance in *Mab*. First, we treated *Mab* cultures in logarithmic phase with amikacin, clarithromycin and cefoxitin, which are commonly prescribed to treat *Mab* infections (67). We found that clarithromycin and cefoxitin alone were ineffective against all of the strains (Fig. 4A). However, amikacin treatment resulted in 10-100-fold decrease in viability of wild-type and complemented strains, but had no effect on Δ*rel_Mab_*.

**Figure 4.**
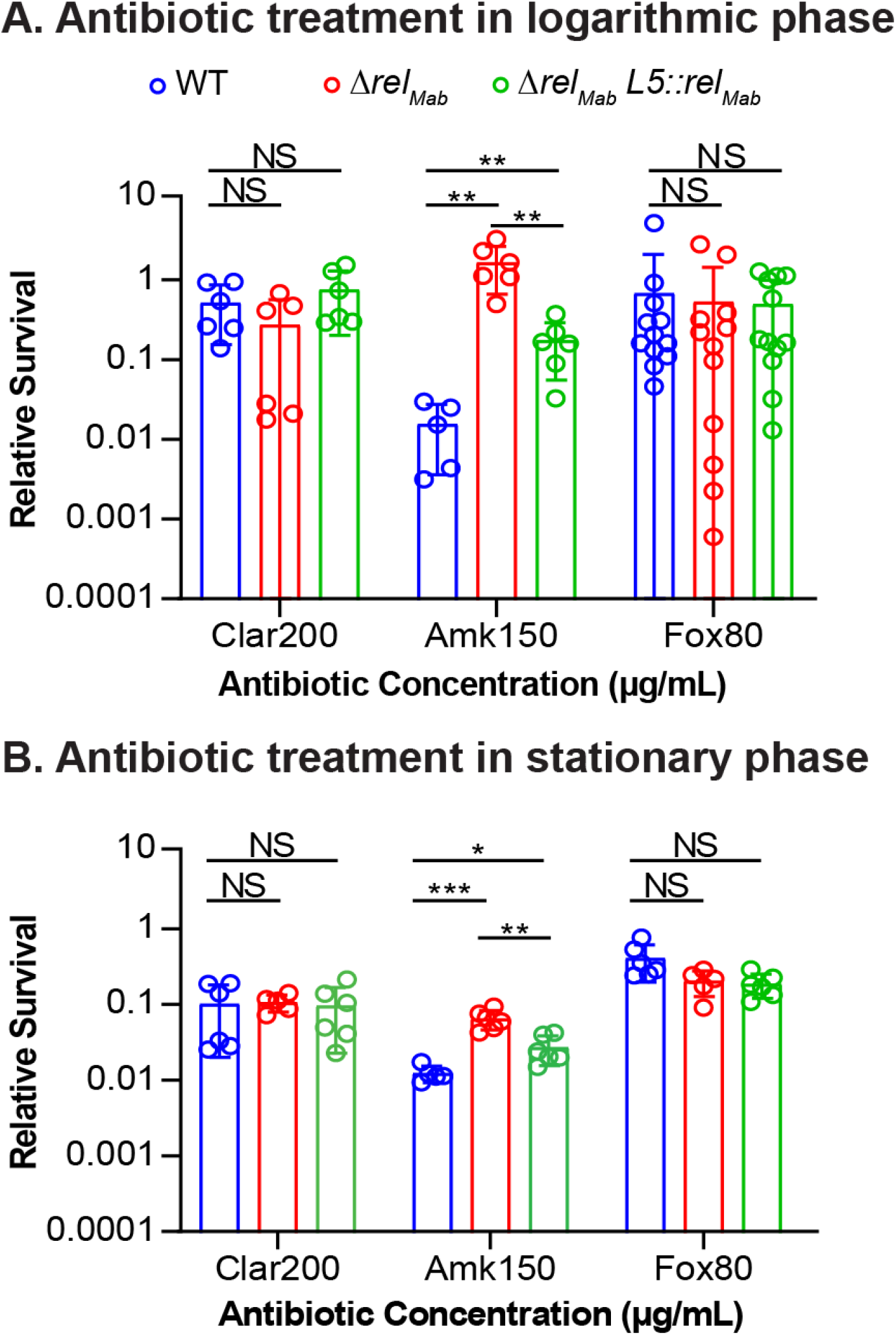
Contribution of *rel_Mab_* to survival in antibiotic treatment. Relative Colony-forming units (CFUs) of strains treated with either 200 μg/mL of clarithromycin (Clar), 150 μg/mL of amikacin (Amk), or 80 μg/mL of cefoxitin (Fox) for either (A) 48 h in logarithmic phase or (B) 72 h in stationary phase. Relative CFUs were calculated by taking the ratio between the CFU value after treatment and the initial CFU value at time zero. The bars represent the mean of 6-9 biological replicates, the individual values are shown by the dots. Error bars represent standard deviation. Logarithmic phase *P*-values: (Amk150) WT vs. Δ*rel_Mab_* = 0.005; WT vs. Δ*rel_Mab_* L5::*rel_Mab_* = 0.01; Δ*rel_Mab_* vs. Δ*rel_Mab_* L5::*rel_Mab_* = 0.004 . Stationary phase *P*-values: (AMK150) WT vs Δ*rel_Mab_* = 0.00017; WT vs. Δ*rel_Mab_* L5::*rel_Mab_* = 0.0217; Δ*rel_Mab_* vs. Δ*rel_Mab_* L5::*rel_Mab_* = 0.0017. Asterisks represent significance as measured by the two-tailed student’s *t*-test; * = *P* ≤ 0.05; ** = *P* ≤ 0.01; *** = *P* ≤ 0.001; n.s. = *P* > 0.05.

Antibiotic tolerance increases in stationary phase in most bacterial species relative to logarithmic phase (5, 68). We repeated the antibiotic survival experiments on cultures in stasis and found that Rel*_Mab_* does not affect tolerance to clarithromycin or cefoxitin (Fig. 4B). Similar to amikacin treatment in growth, Rel*_Mab_* increased susceptibility relative to wild type and the complemented strains in stasis.

A major function of the stringent response in other bacteria is to regulate transcription (4). To determine the effects of Rel*_Mab_* on transcription, we compared the trancriptome of wild-type and Δ*rel_Mab_* in both logarithmic and stationary phases using RNASeq. We found that Rel*_Mab_* represses many more genes than it activates (File. S1).

In logarithmic phase (OD=0.5), when Δ*rel_Mab_* grows more slowly relative to wild type (Fig. 3B), we found 150 genes repressed by *rel_Mab_* in the wild-type strain at least 3-fold, and only 7 genes were activated. The only annotated upregulated genes are an efflux pump (MAB_0677) and a major facilitator superfamily (MFS) transporter (MAB_0069). We found several *mce* family genes that were repressed by *rel_Mab_* (Table S2). Mce proteins are typically lipid transporters, but they also play roles in host cell entry and immune modulation (69). We also found two antibiotic resistance genes that are repressed by *rel_Mab_* in logarithmic phase (Table 1).

**Table 1.**
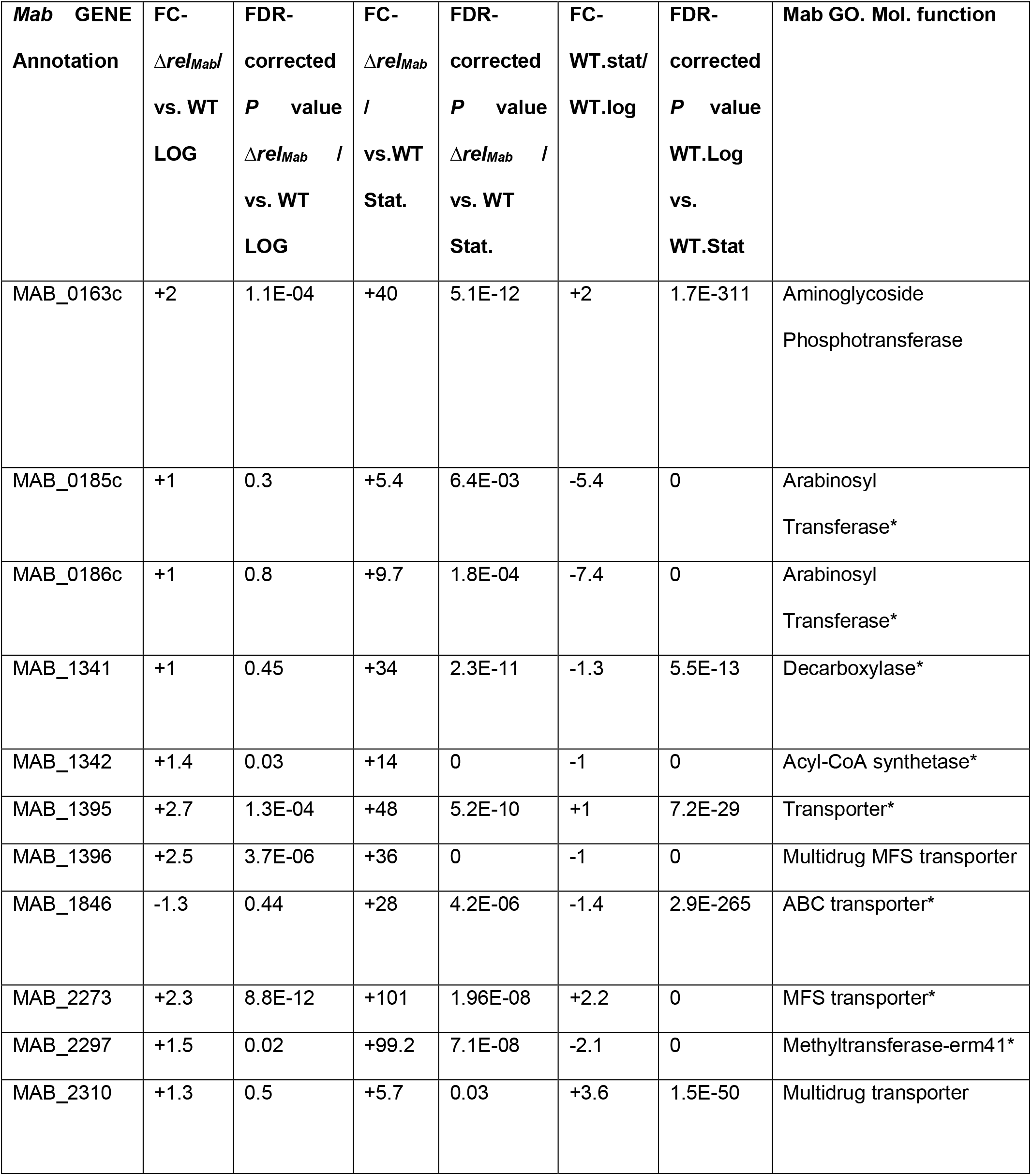
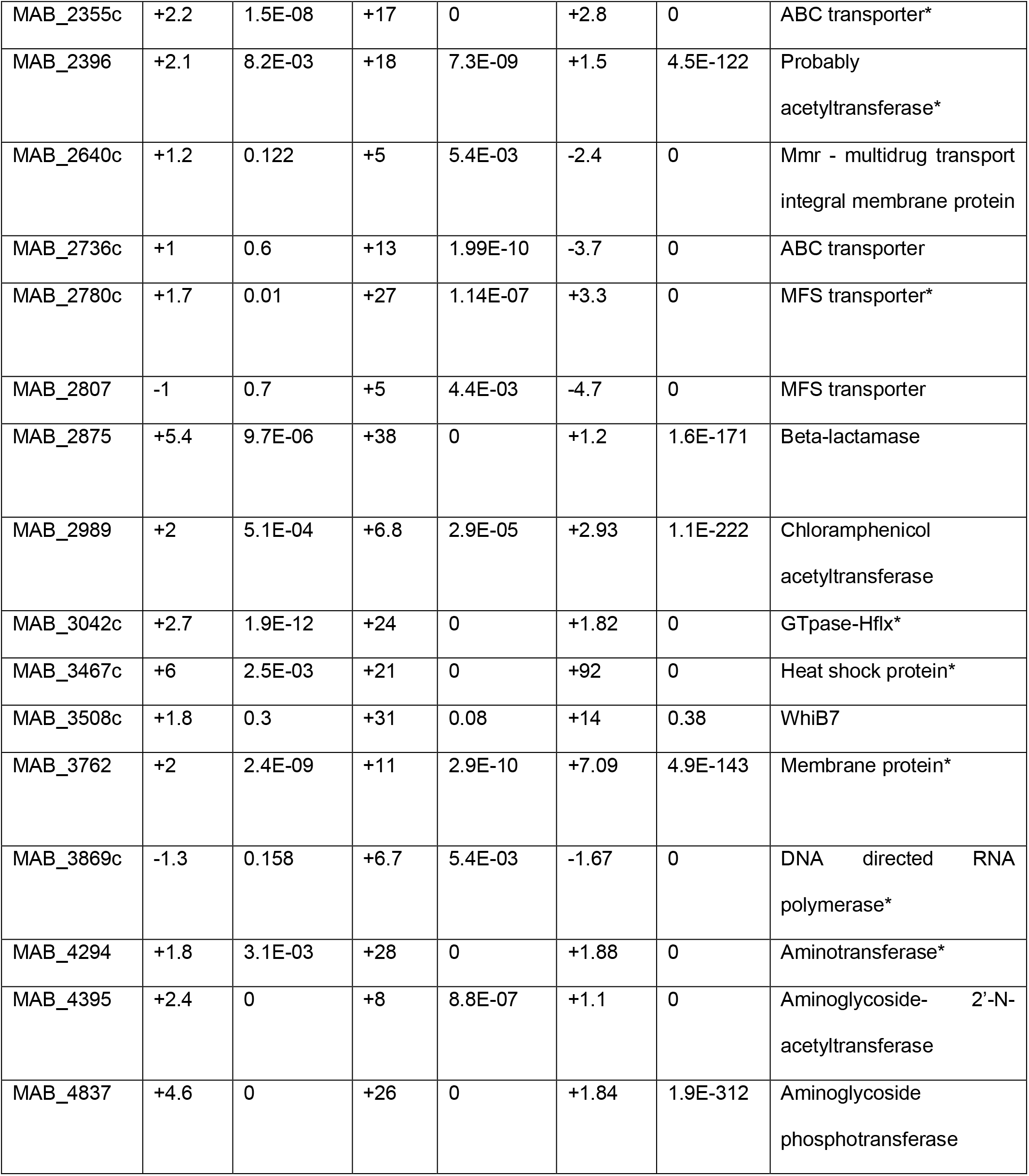
Antibiotic Resistance Genes – (Under whiB7 regulon)*

Even though there was no apparent difference in survival between the wild-type and Δ*rel_Mab_* strains in stationary phase, we observed significant differences in transcription. We isolated RNA from stationary phase cultures that were shaken for 48 hours after being diluted to OD=0.05. We found hundreds of genes that were repressed by *rel_Mab_* in the wild type strain in stationary phase, but none that were activated by *rel_Mab_* 3-fold or more. We found many genes in the WhiB7 regulon (Table 1) which are repressed in stationary phase, though they are mostly unaffected in logarithmic phase. WhiB7 is a transcription factor that activates many antibiotic resistance genes and promotes resistance to many classes of antibiotics in *Mab* (70). It is notable that these antibiotic resistance genes are repressed by *rel_Mab_* in stasis, which would imply that wild type *Mab* would be more susceptible to antibiotics in this condition, which is what we see in amikacin treatment. In the case of clarithromycin and cefoxitin, increased tolerance through downregulation of target expression may counterbalance the repression of the antibiotic resistance genes, resulting in no differences in susceptibility in our assays (Fig. 4).

We also found several cell wall biosynthetic genes that are downregulated by *rel_Mab_* in stationary phase (Table S2). Downregulation of growth factors is typical in stringent responses across many bacterial species (3, 47, 66, 71, 72). We see that *rel_Mab_* downregulates many central metabolism genes in stationary phase (Fig. 5, Table S3), which shows that when Rel is present, it contributes to the decrease in central carbon metabolism during stress. However, it is notable that not all the genes in a given pathway are downregulated equally. We hypothesize that this uneven regulation of certain pathways may allow certain metabolites to accumulate and be re-directed to other pathways. In a metabolic pathway where most of the enzymes are downregulated, we expect that the product of a single gene that is not downregulated will accumulate. We note that many of the metabolites we expect to accumulate converge on the NAD synthesis pathway. In addition, none of the genes in the NAD synthesis pathway are downregulated by *rel_Mab_*, which implies that continued metabolism of NAD, which is a critical cofactor in many pathways, may be important in stationary phase.

**Figure 5.**
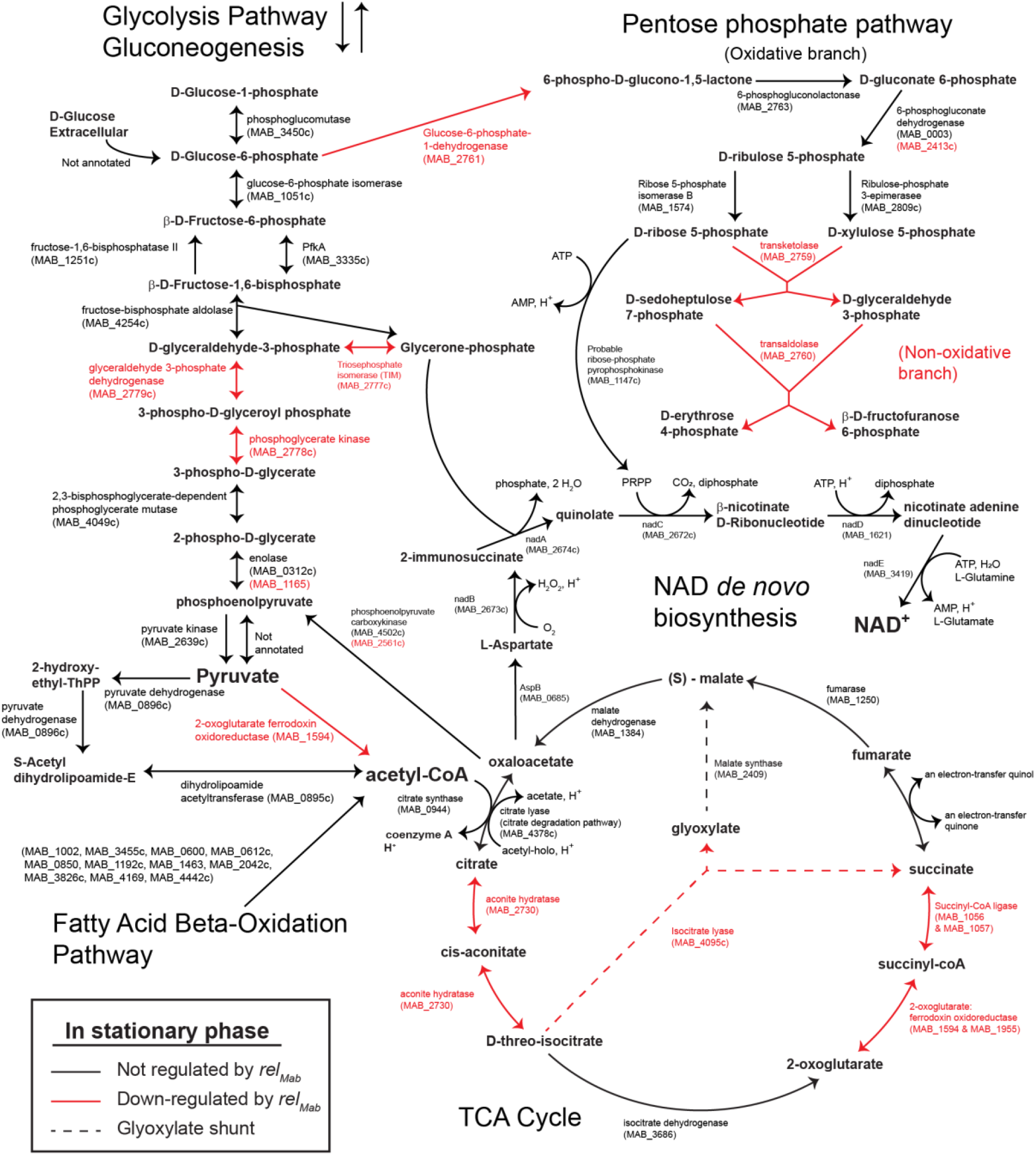
Repression of central metabolic genes by Rel*_Mab_* during stationary phase. Genes in red are downregulated by Rel*_Mab_* at least 3-fold in stationary phase, *P* < 0.05. Genes in black are not significantly regulated by Rel*_Mab_* in stationary phase. See Table S4 for data.

From our preliminary analysis, it is clear that Rel*_Mab_* helps regulate growth and central metabolism, and affects expression of antibiotic resistance genes; however, it does not seem to upregulate specific stress responses in the conditions tested.

## Discussion

*Mtb, Msmeg* and *Mab* all have a stringent response, but they have evolved use of different sets of genes to synthesize the same alarmone. The *Mab* genome encodes only one annotated RSH gene, *rel_Mab_* (MAB_2876), and no homologs of SAS or SAH genes. *rel*_Mab_ is homologous to RSH genes in *Mtb* and *Msmeg*. We found that the Δ*rel_Mab_* strain could still produce (p)ppGpp, consistent with the presence of an unannotated (p)ppGpp synthase. *Mab* does not have a homolog of the SAS-RNaseHII *relZ*, which has been characterized in *Msmeg* (57, 59, 62). Despite this, *Msmeg* and *Mab* are similar in having significant (p)ppGpp synthesis in the absence of the RSH gene (Fig. 1) (53), and in exhibiting greater survival with some antibiotics (Fig. 4) (61, 68). *Mab* does have a homolog of the PNPase which has been shown to be capable of (p)ppGpp synthesis in *Mtb* (MAB_3106c) (56). MAB_3106c is transcriptionally upgregulated 4-fold in the Δ*rel* strain compared to wild-type in stationary phase, indicating that the presence of Rel may decrease the need for transcription of this gene. In the wild-type strain, this gene is also upregulated ~7 fold in stationary phase compared in log. phase, indicating that this gene is used in stress. We therefore propose MAB_3106c as a possible additional (p)ppGpp synthase.

Our results show that the stringent factor Rel*_Mab_* does not promote survival during *in vitro* stress, though it does promote growth during logarithmic phase (Fig. 2, 3A). Our transcriptomics analysis corroborate these results, as they indicate that Rel*_Mab_* does not upregulate stress response genes, but does alter transcription of growth metabolism genes (Table S2, Supplemental file 1). It is possible that increases in (p)ppGpp stimulate transcription of stress response genes in *Mab*, as is seen in other species (3, 52, 73, 74) – our experiments did not test this because in our assays Δ*rel_Mab_* synthesizes comparable amounts of (p)ppGpp as wild-type (Fig. 1). The unannotated ppGpp synthase may contribute to stress response signaling. Our transcriptional data show that Rel*_Mab_* is involved in downregulating metabolism for growth arrest (Fig. 5, S2) even though (p)ppGpp levels are still present in the Δ*rel_Mab_* strain. There is precedent for metabolic genes being regulated separately from stress response genes in the stringent response. In *E. coli*, some metabolic genes are repressed at a lower (p)ppGpp level than is required to induce a regulon of stress response genes (75). (p)ppGpp likely does not directly bind RNAP in mycobacteria as it does in *E. coli* (76), and it is currently unknown how it exerts its effects on transcription. Further knowledge of the functioning of the mycobacterial stringent response will be required before we can speculate about the mechanism by which (p)ppGpp could differentially affect transcription of metabolic and stress response genes in *Mab*.

*Mab* is notorious for having resistance to many clinical antibiotics and expressing many antibiotic resistance genes (70, 77, 78), which is why it is problematic to treat infections in cystic fibrosis patients (79). We observed in our transcriptional data that Rel*_Mab_* downregulated numerous antibiotic resistance genes in stationary phase (Table 1). In other species RSH proteins promote antibiotic tolerance (5, 11, 66) and sometimes also increase expression of antibiotic resistance genes (80, 81). Studies are ongoing to find drugs that would inhibit (p)ppGpp synthesis by RSH proteins (11, 13), as such drugs are expected to increase susceptibility to clinically available antibiotics. Our results indicate that Rel*_Mab_* inhibitors, should they become available, are unlikely to help treat *Mab* infections. Instead, efforts should focus on identifying the other (p)ppGpp synthetase(s) in *Mab* so that it can be explored as a drug target.

## Materials and Methods

### Construction of strains

Primers 1233 – 1238 (Table S3) were used to amplify a 502 bp segment upstream of *Mab rel* which included the start codon, a 448 bp segment downstream of Mab *rel* which included the stop codon, and a 788 bp ZeoR cassette. All 3 segments were stitched together by PCR to form the Δ*rel*::zeoR double stranded recombineering knockout construct. Δ*rel_Mab_* was generated through double stranded recombineering, as previously described (82) (Fig. S1C). Colonies from the transformation of the Δ*rel*::zeoR construct were PCR checked by using primers 1424-761, 1235-1236, and 762-1425 (Fig. S1A). To make the complemented strain, the *rel_Mab_* gene was amplified through PCR using primers 1329-1330 and inserted into pKK216 (83) with NdeI and HindIII. This new plasmid, pCB1248, was transformed into the Δ*rel*_Mab_ mutant strain in order to create the complementation strain, Δ*rel*_Mab_ *L5::rel_Mab_*, in which *rel_Mab_* expression is driven by a constitutive promoter (BN17, Fig.S1B).

### Media and culture conditions

All *M. abscessus* ATCC 19977 wild-type and mutant cultures were started in 7H9 (Becton, Dickinson, Franklin Lakes, NJ) medium with 5 g/liter bovine serum albumin, 2 g/liter dextrose, 0.85 g/liter NaCl, 0.003 g/liter catalase, 0.2% glycerol, and 0.05% Tween 80 and shaken overnight at 37° C until cultures entered logarithmic phase. For starvation and other specific assays, Hartmans-de Bont (HdB) minimal medium was made as described previously (84). Cultures were inoculated to OD_600_ 0.05, unless otherwise stated. All CFU time points were plated on Luria-Bertani (LB) agar and placed in 37° C incubator for 4 days.

### *Mab* (p)ppGpp extraction and detection

*Mab* strains were grown until logarithmic phase (OD_600_ 0.5) in homemade 7H9 medium. To maximize ^32^P-labeling, 7H9 medium with 1/25 of normal phosphate levels was used (46). Logarithmic phase *Mab* cells were pelleted and resuspended in 1mL of low phosphate 7H9, followed by the addition of ^32^P-labeled orthophosphoric acid to a final concentration of 100 μCi/mL. ^32^P-labeled cells were then incubated at 37° C and shaken at 200 rpm for 4 h. Following incubation, cells were pelleted and resuspended with 100 μL of TBST (Tris-buffer saline pH 8, with 0.05% Tyloxapol), and treated with 1 mg/mL of lysozyme on ice for 20 minutes. Equal volume 2 M Formic acid was then added to each sample, followed by a 5 minute centrifugation at max speed. Supernatant was then collected for each sample and stored in −20° C. For starvation cultures, a second set of log phase *Mab* strains (OD_600_ 0.5) were washed and resuspended in TBST. Addition of ^32^P-labeled orthophosphoric acid, and (p)ppGpp extraction was performed similarly as logarithmic phase cultures.. Following extraction, 20 μL of extract from all cultures were spotted on PEI-cellulose TLC plates and developed in 1.5 M Potassium monophosphate buffer (pH 3.4). The plates were then air dried and placed on a phosphor screen (Molecular dynamics) overnight. The phosphor screens were scanned with a Storm 860 scanner (Amersham Biosciences) and images were analyzed with ImageQuant software (Molecular Dynamics).

### Δ*rel_Mab_* stress assays

For all stress assays, strains were prepared and grown into logarithmic phase. Unless otherwise stated, cultures for stress assays were done in 24-well plates and shaken at 130 rpm at 37° C. For carbon starvation, strains were inoculated in 30 mL inkwells in HdB minimal media with no glycerol, and with Tyloxapol as a detergent. For acid stress, strains were inoculated in 7H9 medium pH 4. For osmotic stress, strains were inoculated in LB broth medium with 1M salt (ACS Sodium Chloride, VWR Chemicals BDH). For oxidative stress, all strains were inoculated in complete HdB minimal medium, which does not contain catalase, and strains were exposed to different concentrations of tert-Butyl Hydroperoxide (Alfa Aesar). CFU time points were taken upon inoculation, at 1, 3, and 24 h post-inoculation.

### Δ*rel_Mab_* growth curve and stationary phase survival

Log-phase cultures of all strains were inoculated to OD 0.05 in 30 mL inkwells in 7H9 media. Cultures were then placed in shaking incubator at 37° C and 130 rpm. CFU time points were then taken throughout a 12-hour period. For stationary phase survival, a second set of cultures were grown into stationary phase up to 48 h. Initial CFU time-point was taken at 48 h after dilution of logarithmic phase samples to OD_600_ 0.05, with subsequent time points taken at 5, 6, 7, 8, 9, and 10 d.

### Antibiotic assays

For the logarithmic phase experiments, strains were kept in logarithmic phase in 7H9 for *t*~24 h and diluted to OD_600_ 0.05 and treated with either 150 μg/mL of amikacin, 200 μg/mL clarythromycin, or 80 μg/mL of cefoxitin. These antibiotic concentrations were chosen because they were shown in a previous study to cause killing of *Mab* in certain conditions (9). CFUs were measured upon treatment (*t*=0) and 48 h after treatment (*t*=48). For stationary phase, logarithmic phase cells at OD_600_ 0.05 were shaken for 48 h and then treated as above. CFUs were measured upon treatment and 72 h after treatment.

### RNA isolation, library preparation and data analysis

RNA from three biological replicates of each strain and condition was isolated as previously described (85) with some modifications. After growth for *t*~24 hours in either logrithmic or stationary phase, cells were transferred to 15 mL conical tubes and centrifuged at 4° C for 3 min at 1467 x g. Cell pellets were immediately resuspended in 750 μl of TriZol (Invitrogen) and lysed by bead beating. RNA was purified according to protocol with the Zymogen Direct-zol RNA Miniprep Plus (cat. No 2070). RNA was processed for Illumina sequencing using the TRuSeq Total RNA Library Prep from Illumina, with bacterial rRNA removal probes provided separately by Illumina. Sequencing was performed using Illumina NovaSeq at the North Texas Genome Center at the University of Texas in Arlington.

Between 50-300 million pair-end reads per library were mapped to the *M. abscessus subs. abscessus* strain ATCC 19977 published genome using CLC Genomic Workbench software (Qiagen). To minimize the skewing effect that certain PCR jackpots had on the data, we adjusted the number of reads mapped from each library so the median reads per gene were equivalent within an experiment. In the logarithmic phase samples, the median reads per gene was ~600. In the stationary phase samples, the median reads per gene was ~100. After normalization, the Reads Per Kilobase Million (RPKM) values were determined for each gene, and the weighted proportion fold change of RPKM between wild type and Δ*rel_Mab_* for each condition were calculated by CLC Workbench. The Baggerley’s test was used to generate a false discovery rate corrected P-value. We used an arbitrary cut-off of 3-fold change with a false-discovery rate corrected *P* ≤ 0.05 to identify significantly differentially regulated genes between wild type and Δ*rel_Mab_* in each condition. Because the median reads per gene for logarithmic phase samples was 6 times higher than for stationary phase samples, we linearly scaled the fold-change values when comparing logarithmic to stationary phase data to normalize for this difference in read depth.

### Real-time PCR (Supplemental data)

For RNA-seq validation, real-time PCR (RT-PCR) was performed with the Kapa Biosystems Sybr Fast One-Step qRT-PCR kit. RNA was extracted in triplicates from each *Mab* strain as previously described. Primers for reverse-transcription (RT) were designed for each gene of interest by using Primer3 (Table S3). Each 20 μL RT reaction mixture contained 10 μL One-step SYBR green Master Mix, 1 μL of each primer, 4.5 μL nuclease-free water, 0.5 μL of qScript One-step RT, and 3 μL of RNA. RT-PCR was done by using Bio-Rad CFX Connect Real-time system. The relative target levels (fold change) were calculated using the ΔΔCt method (86), with normalization of RNA targets to TetR transcription factor gene (MAB_1638).

## Acknowledgements

This work was funded by a Pilot and Feasibility Award from the Cystic Fibrosis Foundation to CCB and by funding from the National Institutes of Health (grant AI1468269 and GM143053 to JMB).

## References

1. Dahl JL, Kraus CN, Boshoff HIM, Doan B, Foley K, Avarbock D, Kaplan G, Mizrahi V, Rubin H, Barry CE. 2003. The role of RelMtb-mediated adaptation to stationary phase in long-term persistence of Mycobacterium tuberculosis in mice. Proceedings of the National Academy of Sciences 100:10026–10031.

2. Traxler MF, Summers SM, Nguyen H-T, Zacharia VM, Hightower GA, Smith JT, Conway T. 2008. The global, ppGpp-mediated stringent response to amino acid starvation in Escherichia coli. Molecular Microbiology 68:1128–1148.

3. Boutte CC, Crosson S. 2011. The complex logic of stringent response regulation in Caulobacter crescentus: starvation signalling in an oligotrophic environment. Molecular Microbiology 80:695–714.

4. Hauryliuk V, Atkinson GC, Murakami KS, Tenson T, Gerdes K. 2015. Recent functional insights into the role of (p)ppGpp in bacterial physiology. Nat Rev Microbiol 13:298–309.

5. Harms A, Fino C, Sørensen MA, Semsey S, Gerdes K. 2017. Prophages and Growth Dynamics Confound Experimental Results with Antibiotic-Tolerant Persister Cells. mBio 8.

6. Johansen MD, Herrmann J-L, Kremer L. 2020. Non-tuberculous mycobacteria and the rise of Mycobacterium abscessus. Nat Rev Microbiol https://doi.org/10.1038/s41579-020-0331-1.

7. Nessar R, Cambau E, Reyrat JM, Murray A, Gicquel B. 2012. Mycobacterium abscessus: a new antibiotic nightmare. Journal of Antimicrobial Chemotherapy 67:810–818.

8. Clary G, Sasindran SJ, Nesbitt N, Mason L, Cole S, Azad A, McCoy K, Schlesinger LS, Hall-Stoodley L. 2018. *Mycobacterium abscessus* Smooth and Rough Morphotypes Form Antimicrobial-Tolerant Biofilm Phenotypes but Are Killed by Acetic Acid. Antimicrobial Agents and Chemotherapy 62.

9. Hunt-Serracin AC, Parks BJ, Boll J, Boutte C. 2019. Biofilm-associated Mycobacterium abscessus cells have altered antibiotic tolerance and surface glycolipids in Artificial Cystic Fibrosis Sputum Media. Antimicrobial Agents and Chemotherapy AAC.02488-18.

10. Wexselblatt E, Oppenheimer-Shaanan Y, Kaspy I, London N, Schueler-Furman O, Yavin E, Glaser G, Katzhendler J, Ben-Yehuda S. 2012. Relacin, a Novel Antibacterial Agent Targeting the Stringent Response. PLOS Pathogens 8:e1002925.

11. Dutta NK, Klinkenberg LG, Vazquez M-J, Segura-Carro D, Colmenarejo G, Ramon F, Rodriguez-Miquel B, Mata-Cantero L, Porras-De Francisco E, Chuang Y-M, Rubin H, Lee JJ, Eoh H, Bader JS, Perez-Herran E, Mendoza-Losana A, Karakousis PC. 2019. Inhibiting the stringent response blocks *Mycobacterium tuberculosis* entry into quiescence and reduces persistence. Science Advances 5:eaav2104.

12. Hobbs JK, Boraston AB. 2019. (p)ppGpp and the Stringent Response: An Emerging Threat to Antibiotic Therapy. ACS Infect Dis 5:1505–1517.

13. Pacios O, Blasco L, Bleriot I, Fernandez-Garcia L, Ambroa A, López M, Bou G, Cantón R, Garcia-Contreras R, Wood TK, Tomás M. 2020. (p)ppGpp and Its Role in Bacterial Persistence: New Challenges. Antimicrob Agents Chemother 64:e01283–20, /aac/64/10/AAC.01283-20.atom.

14. Barker MM, Gaal T, Josaitis CA, Gourse RL. 2001. Mechanism of regulation of transcription initiation by ppGpp. I. Effects of ppGpp on transcription initiation in vivo and in vitro1 1 Edited by R. Ebright. Journal of Molecular Biology 305:673–688.

15. Barker MM, Gaal T, Gourse RL. 2001. Mechanism of regulation of transcription initiation by ppGpp. II. Models for positive control based on properties of RNAP mutants and competition for RNAP11Edited by R. Ebright. Journal of Molecular Biology 305:689–702.

16. Krásný L, Gourse RL. 2004. An alternative strategy for bacterial ribosome synthesis: Bacillus subtilis rRNA transcription regulation. EMBO J 23:4473–4483.

17. Krásný L, Tišerová H, Jonák J, Rejman D, Šanderová H. 2008. The identity of the transcription +1 position is crucial for changes in gene expression in response to amino acid starvation in Bacillus subtilis. Molecular Microbiology 69:42–54.

18. Maciąg M, Kochanowska M, Łyżeń R, Węgrzyn G, Szalewska-Pałasz A. 2010. ppGpp inhibits the activity of Escherichia coli DnaG primase. Plasmid 63:61–67.

19. Kraemer JA, Sanderlin AG, Laub MT. 2019. The Stringent Response Inhibits DNA Replication Initiation in *E. coli* by Modulating Supercoiling of *oriC*. mBio 10:e01330–19, /mbio/10/4/mBio.01330-19.atom.

20. Kriel A, Bittner AN, Kim SH, Liu K, Tehranchi AK, Zou WY, Rendon S, Chen R, Tu BP, Wang JD. 2012. Direct Regulation of GTP Homeostasis by (p)ppGpp: A Critical Component of Viability and Stress Resistance. Molecular Cell 48:231–241.

21. Wang B, Dai P, Ding D, Del Rosario A, Grant RA, Pentelute BL, Laub MT. 2019. Affinity-based capture and identification of protein effectors of the growth regulator ppGpp. 2. Nature Chemical Biology 15:141–150.

22. Wang B, Grant RA, Laub MT. 2020. ppGpp Coordinates Nucleotide and Amino-Acid Synthesis in E. coli During Starvation. Molecular Cell S1097276520305487.

23. Fan H, Hahm J, Diggs S, Perry JJP, Blaha G. 2015. Structural and Functional Analysis of BipA, a Regulator of Virulence in Enteropathogenic *Escherichia coli*. J Biol Chem 290:20856–20864.

24. Corrigan RM, Bellows LE, Wood A, Gründling A. 2016. ppGpp negatively impacts ribosome assembly affecting growth and antimicrobial tolerance in Gram-positive bacteria. Proc Natl Acad Sci USA 113:E1710–E1719.

25. Rojas A-M, Ehrenberg M, Andersson SGE, Kurland CG. 1984. ppGpp inhibition of elongation factors Tu, G and Ts during polypeptide synthesis. Mol Gen Genet 197:36–45.

26. Milon P, Tischenko E, Tomsic J, Caserta E, Folkers G, La Teana A, Rodnina MV, Pon CL, Boelens R, Gualerzi CO. 2006. The nucleotide-binding site of bacterial translation initiation factor 2 (IF2) as a metabolic sensor. Proceedings of the National Academy of Sciences 103:13962–13967.

27. Mitkevich VA, Ermakov A, Kulikova AA, Tankov S, Shyp V, Soosaar A, Tenson T, Makarov AA, Ehrenberg M, Hauryliuk V. 2010. Thermodynamic Characterization of ppGpp Binding to EF-G or IF2 and of Initiator tRNA Binding to Free IF2 in the Presence of GDP, GTP, or ppGpp. Journal of Molecular Biology 402:838–846.

28. Atkinson GC, Tenson T, Hauryliuk V. 2011. The RelA/SpoT Homolog (RSH) Superfamily: Distribution and Functional Evolution of ppGpp Synthetases and Hydrolases across the Tree of Life. PLoS ONE 6:e23479.

29. Haseltine WA, Block R. 1973. Synthesis of Guanosine Tetra- and Pentaphosphate Requires the Presence of a Codon-Specific, Uncharged Transfer Ribonucleic Acid in the Acceptor Site of Ribosomes. Proc Natl Acad Sci U S A 70:1564–1568.

30. Ronneau S, Hallez R. 2019. Make and break the alarmone: regulation of (p)ppGpp synthetase/hydrolase enzymes in bacteria. FEMS Microbiology Reviews 43:389–400.

31. Battesti A, Bouveret E. 2006. Acyl carrier protein/SpoT interaction, the switch linking SpoT-dependent stress response to fatty acid metabolism. Molecular Microbiology 62:1048–1063.

32. Brown DR, Barton G, Pan Z, Buck M, Wigneshweraraj S. 2014. Nitrogen stress response and stringent response are coupled in Escherichia coli. Nat Commun 5:4115.

33. Ronneau S, Petit K, De Bolle X, Hallez R. 2016. Phosphotransferase-dependent accumulation of (p)ppGpp in response to glutamine deprivation in Caulobacter crescentus. Nat Commun 7:11423.

34. Germain E, Guiraud P, Byrne D, Douzi B, Djendli M, Maisonneuve E. 2019. YtfK activates the stringent response by triggering the alarmone synthetase SpoT in Escherichia coli. Nat Commun 10:5763.

35. Nanamiya H, Kasai K, Nozawa A, Yun C-S, Narisawa T, Murakami K, Natori Y, Kawamura F, Tozawa Y. 2007. Identification and functional analysis of novel (p)ppGpp synthetase genes in Bacillus subtilis: Novel (p)ppGpp synthetase genes in B. subtilis. Molecular Microbiology 67:291–304.

36. Ruwe M, Kalinowski J, Persicke M. 2017. Identification and Functional Characterization of Small Alarmone Synthetases in Corynebacterium glutamicum. Front Microbiol 8:1601.

37. Ruwe M, Rückert C, Kalinowski J, Persicke M. 2018. Functional Characterization of a Small Alarmone Hydrolase in Corynebacterium glutamicum. Front Microbiol 9:916.

38. Yang N, Xie S, Tang N-Y, Choi MY, Wang Y, Watt RM. 2019. The Ps and Qs of alarmone synthesis in Staphylococcus aureus. PLoS ONE 14:e0213630.

39. Jimmy S, Saha CK, Kurata T, Stavropoulos C, Oliveira SRA, Koh A, Cepauskas A, Takada H, Rejman D, Tenson T, Strahl H, Garcia-Pino A, Hauryliuk V, Atkinson GC. 2020. A widespread toxin-antitoxin system exploiting growth control via alarmone signaling. Proc Natl Acad Sci USA 117:10500–10510.

40. Jones GH. 1994. Purification and properties of ATP:GTP 3’-pyrophosphotransferase (guanosine pentaphosphate synthetase) from Streptomyces antibioticus. J Bacteriol 176:1475–1481.

41. Jones GH. 1994. Activation of ATP:GTP 3’-pyrophosphotransferase (guanosine pentaphosphate synthetase) in Streptomyces antibioticus. J Bacteriol 176:1482–1487.

42. Jones GH, Bibb MJ. 1996. Guanosine pentaphosphate synthetase from Streptomyces antibioticus is also a polynucleotide phosphorylase. Journal of bacteriology 178:4281–4288.

43. Jones GH. 2018. Novel Aspects of Polynucleotide Phosphorylase Function in Streptomyces. Antibiotics (Basel) 7:25.

44. Schofield WB, Zimmermann-Kogadeeva M, Zimmermann M, Barry NA, Goodman AL. 2018. The Stringent Response Determines the Ability of a Commensal Bacterium to Survive Starvation and to Persist in the Gut. Cell Host & Microbe 24:120–132.e6.

45. Cashel M, Gallant J. 1969. Two Compounds implicated in the Function of the RC Gene of Escherichia coli. Nature 221:838.

46. Primm TP, Andersen SJ, Mizrahi V, Avarbock D, Rubin H, Barry CE. 2000. The Stringent Response of Mycobacterium tuberculosis Is Required for Long-Term Survival. Journal of Bacteriology 182:4889–4898.

47. Eymann C, Homuth G, Scharf C, Hecker M. 2002. Bacillus subtilis functional genomics: global characterization of the stringent response by proteome and transcriptome analysis. JB 184:2500–2520.

48. Potrykus K, Murphy H, Philippe N, Cashel M. 2011. ppGpp is the major source of growth rate control in E. coli. Environmental Microbiology 13:563–575.

49. Weiss LA, Stallings CL. 2013. Essential Roles for Mycobacterium tuberculosis Rel beyond the Production of (p)ppGpp. Journal of Bacteriology 195:5629–5638.

50. Pulschen AA, Sastre DE, Machinandiarena F, Asis AC, Albanesi D, Mendoza D de, Gueiros-Filho FJ. 2017. The stringent response plays a key role in Bacillus subtilis survival of fatty acid starvation. Molecular Microbiology 103:698–712.

51. Murch AL, Skipp PJ, Roach PL, Oyston PCF. 2017. Whole genome transcriptomics reveals global effects including up-regulation of Francisella pathogenicity island gene expression during active stringent response in the highly virulent Francisella tularensis subsp. tularensis SCHU S4. Microbiology 163:1664–1679.

52. Schäfer H, Beckert B, Frese CK, Steinchen W, Nuss AM, Beckstette M, Hantke I, Driller K, Sudzinová P, Krásný L, Kaever V, Dersch P, Bange G, Wilson DN, Turgay K. 2020. The alarmones (p)ppGpp are part of the heat shock response of Bacillus subtilis. PLoS Genet 16:e1008275.

53. Stallings CL, Stephanou NC, Chu L, Hochschild A, Nickels BE, Glickman MS. 2009. CarD Is an Essential Regulator of rRNA Transcription Required for Mycobacterium tuberculosis Persistence. Cell 138:146–159.

54. Klinkenberg LG, Lee J, Bishai WR, Karakousis PC. 2010. The Stringent Response Is Required for Full Virulence of *Mycobacterium tuberculosis* in Guinea Pigs. J INFECT DIS 202:1397–1404.

55. Prusa J, Zhu DX, Stallings CL. 2018. The stringent response and Mycobacterium tuberculosis pathogenesis. Pathogens and Disease; Oxford 76.

56. Njire M, Wang N, Wang B, Tan Y, Cai X, Liu Y, Mugweru J, Guo J, Hameed HMA, Tan S, Liu J, Yew WW, Nuermberger E, Lamichhane G, Liu J, Zhang T. 2017. Pyrazinoic Acid Inhibits a Bifunctional Enzyme in Mycobacterium tuberculosis. Antimicrob Agents Chemother 61:e00070–17, e00070-17.

57. Dahl JL, Arora K, Boshoff HI, Whiteford DC, Pacheco SA, Walsh OJ, Lau-Bonilla D, Davis WB, Garza AG. 2005. The relA Homolog of Mycobacterium smegmatis Affects Cell Appearance, Viability, and Gene Expression. Journal of Bacteriology 187:2439–2447.

58. Jain V, Saleem-Batcha R, China A, Chatterji D. 2006. Molecular dissection of the mycobacterial stringent response protein Rel. Protein Science 15:1449–1464.

59. Murdeshwar MS, Chatterji D. 2012. MS_RHII-RSD, a Dual-Function RNase HII-(p)ppGpp Synthetase from Mycobacterium smegmatis. Journal of Bacteriology 194:4003–4014.

60. Ojha AK, Mukherjee TK, Chatterji D. 2000. High Intracellular Level of Guanosine Tetraphosphate in Mycobacterium smegmatis Changes the Morphology of the Bacterium. Infect Immun 68:4084–4091.

61. Gupta KR, Kasetty S, Chatterji D. 2015. Novel Functions of (p)ppGpp and Cyclic di-GMP in Mycobacterial Physiology Revealed by Phenotype Microarray Analysis of Wild-Type and Isogenic Strains of Mycobacterium smegmatis. Appl Environ Microbiol 81:2571–2578.

62. Petchiappan A, Naik SY, Chatterji D. 2019. RelZ-mediated stress response in *Mycobacterium smegmatis* : pGpp synthesis and its regulation. J Bacteriol JB.00444-19, jb;JB.00444-19v1.

63. Sugisaki K, Hanawa T, Yonezawa H, Osaki T, Fukutomi T, Kawakami H, Yamamoto T, Kamiya S. 2013. Role of (p)ppGpp in biofilm formation and expression of filamentous structures in Bordetella pertussis. Microbiology 159:1379–1389.

64. Oh YT, Lee K-M, Bari W, Raskin DM, Yoon SS. 2015. (p)ppGpp, a Small Nucleotide Regulator, Directs the Metabolic Fate of Glucose in *Vibrio cholerae*. J Biol Chem 290:13178–13190.

65. Kim J-S, Liu L, Fitzsimmons LF, Wang Y, Crawford MA, Mastrogiovanni M, Trujillo M, Till JKA, Radi R, Dai S, Vázquez-Torres A. 2018. DksA–DnaJ redox interactions provide a signal for the activation of bacterial RNA polymerase. Proceedings of the National Academy of Sciences 115:E11780–E11789.

66. Gaca AO, Kajfasz JK, Miller JH, Liu K, Wang JD, Abranches J, Lemos JA. 2013. Basal Levels of (p)ppGpp in Enterococcus faecalis: the Magic beyond the Stringent Response. mBio 4:e00646–13.

67. Novosad SA, Beekmann SE, Polgreen PM, Mackey K, Winthrop KL. 2016. Treatment of Mycobacterium abscessus Infection. Emerg Infect Dis 22:511–514.

68. Cabral DJ, Wurster JI, Belenky P. 2018. Antibiotic Persistence as a Metabolic Adaptation: Stress, Metabolism, the Host, and New Directions. Pharmaceuticals (Basel) 11.

69. Fenn K, Wong CT, Darbari VC. 2020. Mycobacterium tuberculosis Uses Mce Proteins to Interfere With Host Cell Signaling. Front Mol Biosci 6:149.

70. Hurst-Hess K, Rudra P, Ghosh P. 2017. Mycobacterium abscessus WhiB7 Regulates a Species-Specific Repertoire of Genes To Confer Extreme Antibiotic Resistance. Antimicrobial Agents and Chemotherapy 61.

71. Brockmann-Gretza O, Kalinowski J. 2006. Global gene expression during stringent response in Corynebacterium glutamicum in presence and absence of the rel gene encoding (p)ppGpp synthase. BMC Genomics 7:230.

72. Traxler MF, Summers SM, Nguyen H-T, Zacharia VM, Hightower GA, Smith JT, Conway T. 2008. The global, ppGpp-mediated stringent response to amino acid starvation in Escherichia coli. Molecular Microbiology 68:1128–1148.

73. Vercruysse M, Fauvart M, Jans A, Beullens S, Braeken K, Cloots L, Engelen K, Marchal K, Michiels J. 2011. Stress response regulators identified through genome-wide transcriptome analysis of the (p)ppGpp-dependent response in Rhizobium etli. Genome Biol 12:R17.

74. Yang H, Yu M, Lee JH, Chatnaparat T, Zhao Y. 2020. The stringent response regulator (p) ppGpp mediates virulence gene expression and survival in Erwinia amylovora. BMC Genomics 21:261.

75. Traxler MF, Zacharia VM, Marquardt S, Summers SM, Nguyen H-T, Stark SE, Conway T. 2011. Discretely calibrated regulatory loops controlled by ppGpp partition gene induction across the ‘feast to famine’ gradient in Escherichia coli: Architecture of the stringent response. Molecular Microbiology 79:830–845.

76. Ross W, Vrentas CE, Sanchez-Vazquez P, Gaal T, Gourse RL. The Magic Spot: A ppGpp Binding Site on E. coli RNA Polymerase Responsible for Regulation of Transcription Initiation. Molecular Cell 10.

77. Jayasingam S, Zin T, Ngeow Y. 2017. Antibiotic resistance in Mycobacterium Abscessus and Mycobacterium Fortuitum isolates from Malaysian patients. Int J Mycobacteriol 6:387.

78. Luthra S, Rominski A, Sander P. 2018. The Role of Antibiotic-Target-Modifying and Antibiotic-Modifying Enzymes in Mycobacterium abscessus Drug Resistance. Front Microbiol 9.

79. Bar-On O, Mussaffi H, Mei-Zahav M, Prais D, Steuer G, Stafler P, Hananya S, Blau H. 2015. Increasing nontuberculous mycobacteria infection in cystic fibrosis. Journal of Cystic Fibrosis 14:53–62.

80. Koskiniemi S, Pränting M, Gullberg E, Näsvall J, Andersson DI. 2011. Activation of cryptic aminoglycoside resistance in Salmonella enterica: Activation of cryptic resistance. Molecular Microbiology 80:1464–1478.

81. Aedo S, Tomasz A. 2016. Role of the Stringent Stress Response in the Antibiotic Resistance Phenotype of Methicillin-Resistant Staphylococcus aureus. Antimicrob Agents Chemother 60:2311–2317.

82. van Kessel JC, Hatfull GF. 2007. Recombineering in Mycobacterium tuberculosis. Nat Methods 4:147–152.

83. Kieser KJ, Boutte CC, Kester JC, Baer CE, Barczak AK, Meniche X, Chao MC, Rego EH, Sassetti CM, Fortune SM, Rubin EJ. 2015. Phosphorylation of the Peptidoglycan Synthase PonA1 Governs the Rate of Polar Elongation in Mycobacteria. PLOS Pathogens 11:e1005010.

84. Hartmans S, de Bont JAM, Stackebrandt E. 2006. The Genus Mycobacterium--Nonmedical, p. 889–918. In Dworkin, M, Falkow, S, Rosenberg, E, Schleifer, K-H, Stackebrandt, E (eds.), The Prokaryotes. Springer New York, New York, NY.

85. Shell SS, Prestwich EG, Baek S-H, Shah RR, Sassetti CM, Dedon PC, Fortune SM. 2013. DNA Methylation Impacts Gene Expression and Ensures Hypoxic Survival of Mycobacterium tuberculosis. PLoS Pathog 9:e1003419.

86. Livak KJ, Schmittgen TD. 2001. Analysis of Relative Gene Expression Data Using Real-Time Quantitative PCR and the 2-ΔΔCT Method. Methods 25:402–408.

